# Host-parasite tissue adhesion by a secreted type of β-1,4-glucanase in the parasitic plant *Phtheirospermum japonicum*

**DOI:** 10.1101/2020.03.29.014886

**Authors:** Ken-ichi Kurotani, Takanori Wakatake, Yasunori Ichihashi, Koji Okayasu, Yu Sawai, Satoshi Ogawa, Takamasa Suzuki, Ken Shirasu, Michitaka Notaguchi

## Abstract

Tissue adhesion between plant species occurs both naturally and artificially. Parasitic plants establish symbiotic relationship with host plants by adhering tissues at roots or stems. Plant grafting, on the other hand, is a widely used technique in agriculture to adhere tissues of two stems. While compatibility of tissue adhesion in plant grafting is often limited within close relatives, parasitic plants exhibit much wider compatibilities. For example, the Orobanchaceae parasitic plant *Striga hermonthica* is able to infect Poaceae crop plants, causing a serious agricultural loss. Here we found that the model Orobanchaceae parasite plant *Phtheirospermum japonicum* can be grafted on to interfamily species, such as *Arabidopsis*, a Brassicaceae plant. To understand molecular basis of tissue adhesion between distant plant species, we conducted comparative transcriptome analyses on both infection and grafting by *P. japonicum* on *Arabidopsis*. Through gene clustering, we identified genes upregulated during these tissue adhesion processes, which include cell proliferation- and cell wall modification-related genes. By comparing with a transcriptome dataset of interfamily grafting between *Nicotiana* and *Arabidopsis*, we identified 9 genes commonly induced in tissue adhesion between distant species. Among them, we showed a gene encoding secreted type of β-1,4-glucanase plays an important role for plant parasitism. Our data provide insights into the molecular commonality between parasitism and grafting in plants.

**Significance Statement:** Comprehensive sequential RNA-Seq datasets for parasitic infection of the root and grafting of the stem between *P. japonicum* and *Arabidopsis* revealed that molecular events of parasitism and grafting are substantially different and only share a part of events such as cell proliferation and cell wall modification. This study demonstrated that a secreted type of β-1,4-glucanase gene expressed in cells located at the parasite–host interface as an important factor for parasitism in the Orobanchaceae.

## Introduction

Exceptionally in the autotrophic plant lineage, parasitic plants have evolved the capability to absorb water and nutrients from other plants. This ability relies on a specialized organ called a haustorium, which forms a physical and physiological connection between the parasite and host tissues (1). Plant parasitism has independently evolved in angiosperm lineages at least 12 times and approximately 1% of angiosperms are estimated to be parasitic (2, 3). Among these species, the Orobanchaceae family is the most species-rich and includes the notorious agricultural pests *Striga* spp., *Phelipanche*, and *Orobanche* spp., which threaten world food security (4).

The infection process of parasitic plant of host plant tissues is initiated with physical contact between the parasitic haustorium and the host tissue, followed by adhesion between them. Electron micrographs of the interaction between *Striga* haustorium and a host show that parasitic ingression is accompanied by host cell wall alterations, but not disruption, such as partial wall dissolution and shredding (5). Similarly, *Orobanche* spp. penetrate the host root tissues where pectolytic enzyme activity is evident around haustoria (6). Activities of cell wall-degrading enzymes, such as cellulase and polygalacturonase, are also present in infecting *Phelipanche* tubers (7). In the case of stem parasites, such as dodder (*Cuscuta pentagona*), epidermal cells differentiate into secretory trichomes that excrete a cementing substance predominantly composed of de-esterified pectins, and the cell walls are modified by a cell-wall-loosening complex (8). The parasitic haustorium thus is able to adhere to the host tissues either in roots or in stems.

All the parasitic plants known to date are able to establish vasculature connection to host, which can be considered as “natural grafting.” Especially, one of the interesting characteristics of parasitic plants is their ability to adhere to the apoplastic cell wall matrix of phylogenetically distant plant species of diverse cell wall composition. This is remarkable as “artificial grafting”, in which cut stem tissues are assembled to unite, often causes incompatibility among interfamily species (9). In the case of compatible graft combinations, the grafted parts are connected through tissue adhesion. Compressed cell walls in the region of the graft junction have been observed during grafting (10, 11), which indicates that the cell walls between opposing cells at the graft interface adhere followed by vascular reconstruction and tissue union between the grafted organs. The mechanism of how parasitic plants are able to overcome incompatibility in tissue adhesion with a diverse range of host plant species remains unclear.

To understand molecular events during parasite infection, transcriptome analyses have been conducted on several parasitic plants, including species of Orobanchaceae, as well as dodder (12–17). In particular, Yang *et al*. (2014) identified putative parasitism genes that are upregulated during haustorial development following host attachment in three Orobanchaceae parasitic species (12). Among them, genes that encode proteases, cell wall modification enzymes, and extracellular secretion proteins are highly upregulated. Similarly, transcriptome analysis of dodder revealed increased expression of genes encoding cell wall modifying enzymes, such as pectin lyase, pectin methyl esterase, cellulase, and expansins, in the infective stages (13). A transcriptome analysis of *Thesium chinense*, a parasitic Santalaceae plant, also identified highly upregulated genes that encode proteins associated with cell wall organization as a peripheral module in the gene co-expression network during developmental reprogramming of haustorium (18). In addition, upregulation of genes that encode cell wall-modifying enzymes was detected in the transcriptome of host–parasite interface in the model *Triphysaria versicolor*, using laser microdissection (19). These aforementioned results suggest that parasitic plants facilitate cell–cell adhesion at the interface between the haustorium and host through activation of cell wall-modification enzymes.

In this study, we addressed molecular commonality between parasitic infection and artificial grafting by comparing tissue adhesion events between *P. japonicum* and *Arabidopsis*. Although these events occur in different organs, we expected that key common components for tissue adhesion would be found by comparative transcriptomic analyses. In addition, we further compared these datasets with that of interfamily graft of *Nicotiana benthamiana*, which is able to adhere cells with those of plant species from diverse families in grafting (20). We identified nine genes that were commonly upregulated in *P. japonicum* haustorium, *P. japonicum*/*Arabidopsis* and *N. benthamiana*/*Arabidopsis* grafting sites. Among them, we identified a gene encoding β-1,4-glucanase as an important factor in plant parasitism.

## Results

### Tissue adhesion between *P. japonicum* and *Arabidopsis* in parasitism and grafting

A facultative parasitic plant, *P. japonicum*, has been studied previously as a model root parasite that can parasitize *Arabidopsis* (15, 21, 22). The ability to transport materials from *Arabidopsis* to *P. japonicum* can be visualized using a symplasmic tracer dye, carboxyfluorescein (CF) (22) (Fig. 1 *A* and *B*). At the parasitization site in the root, a xylem bridge is formed in the haustorium (Fig. 1*C*), by which the *P. japonicum* tissues invade the host tissues (Fig. 1*D*). We observed the interface of the *P. japonicum* haustorium and *Arabidopsis* root tissues using transmission electron microscopy (Fig. 1 *E–I*). The cells at the tip of the penetrating haustoria adhered closely to the opposing *Arabidopsis* cells where thin cell walls were observed (Fig. 1*E*). Serial sections revealed a decrease in cell wall thickness at the interface between *P. japonicum* and *Arabidopsis* tissues (Fig. 1 *F–I*), which indicated that cell wall digestion occurred at the interface.

**Fig. 1.**
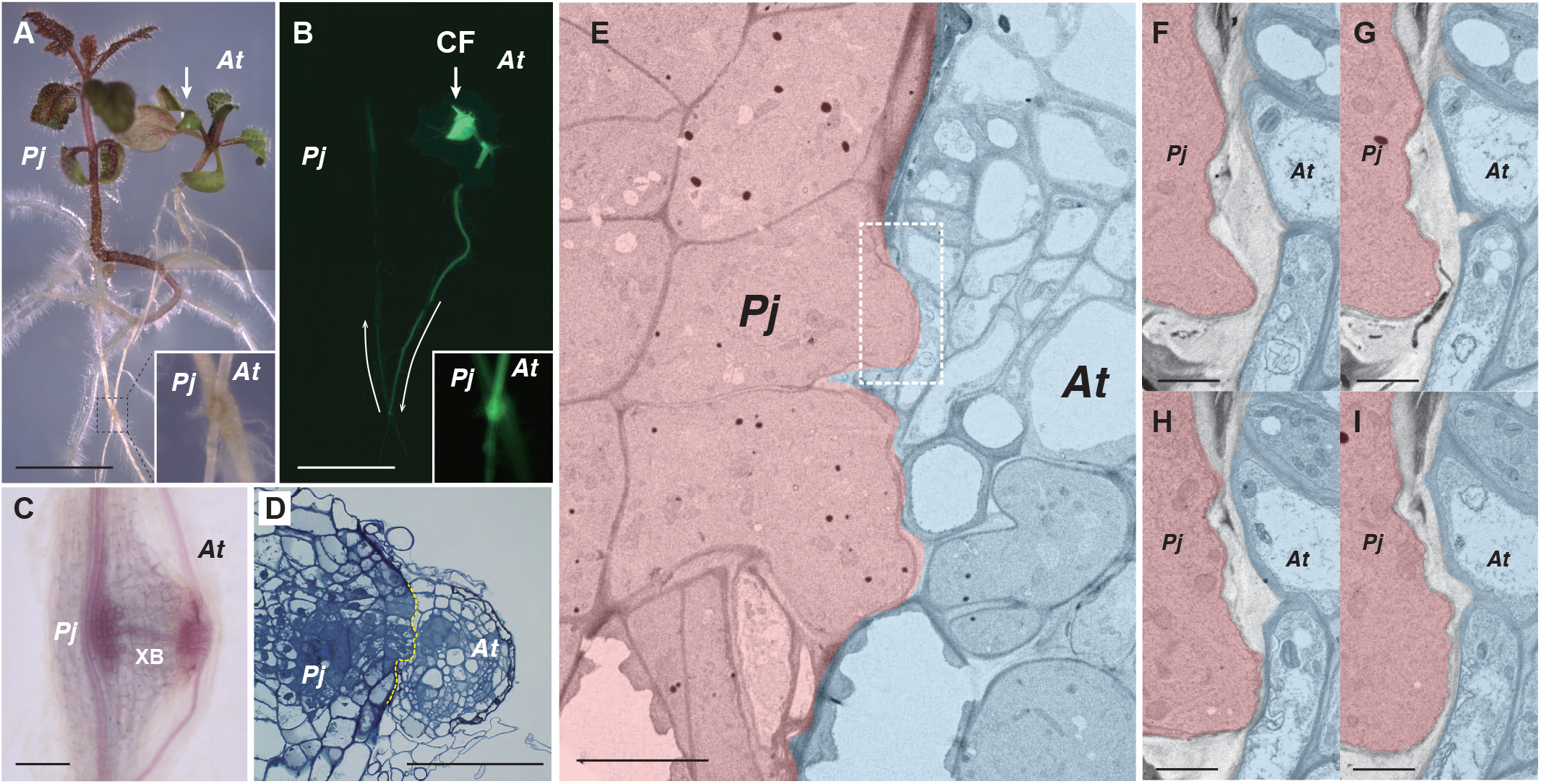
Parasitism of *Phtheirospermum japonicum.* (*A*, *B*) Parasitism between the roots of *P. japonicum* and *Arabidopsis*. The *P. japonicum* root parasitized the *Arabidopsis* root (insets). Transport of a symplasmic tracer dye, carboxyfluorescein (CF, coloured in green), showed establishment of a symplasmic connection between the plants; light micrograph (*A*) and fluorescence image (*B*). Arrows indicate the site where CF dye was applied and the direction of transport. (*C*, *D*) Site where *P. japonicum* parasitized the *Arabidopsis* root. (*C*) Phloroglucinol staining showing xylem bridge formation (XB). (*D*) Cross-section of the parasitization site. The *P. japonicum* tissue invaded the *Arabidopsis* root tissues. Dashed line indicates the interface of parasitism. (*E*) Transmission electron micrograph of the interface between *P. japonicum* (pink) and *Arabidopsis* (blue). Partial tissue adhesion was observed at the interface. The dashed rectangle indicates the area of (*F*–*I*). (*F*–*I*) Serial sections at the interface between *P. japonicum* and *Arabidopsis* cells. The cell wall was partially digested. *Pj*, *P. japonicum*; *At*, *Arabidopsis.* Scale bars, 5 mm (*A*, *B*), 100 μm (*C*, *D*), 10 μm (*E*), and 2 μm (*F*–*I*).

We observed similar thin cell walls at graft boundary between *Arabidopsis* and *Nicotiana* species, which exhibit a capability to adhere their tissue across interfamily species (20, 23) (Fig. 2*A* and *B*). Therefore, we hypothesized that the parasitic plant may also have a wide tissue adhesion capability in artificial grafting. To test this hypothesis, we grafted a stem of *P. japonicum* (as the scion) onto the *Arabidopsis* inflorescence stem. Callus cells proliferated on the cut surface of the scion, similar to haustorial formation. The *P. japonicum* scion was able to establish a graft union with the *Arabidopsis* stock and remained viable for 1 month after grafting (Fig. 2*C*). Given that parasitic Orobanchaceae species have a diverse host range among angiosperms (24), we further tested graft combinations using nine species from seven orders of angiosperms. The grafting capability of *P. japonicum* as the scion using these interfamily species as the stock were clearly observed, except for two Fabaceae species (Fig. 2 *D*–*F*, Table 1). Reciprocally, *P. japonicum* was able to be used as the stock plant for certain plant species (Fig. 2*G*, Table 1). In contrast, when *Lindenbergia philippensis*, which has no parasitic ability among Orobanchaceae (4), was grafted onto *Arabidopsis*, viability of the *L. philippensis* scion was extremely limited compared with the *P. japonicum* scion; 9 graft trials were never successful, although all 9 homografts of *L. philippensis* were successful (Table 1, *SI Appendix*, Fig. S2*C*). When we observed cross-sections of the graft junction of *P. japonicum*/*Arabidopsis* (scion/stock), xylem continuity and apoplastic dye transport were observed (Fig. 2 *H* and *I*). Importantly, establishment of the symplast between *P. japonicum* and *Arabidopsis* was confirmed by using the CF dye (Fig. 2*L*). In summary, these results showed that the root parasite *P. japonicum* is able to achieve tissue adhesion and vasculature connection with members of diverse plant families in both parasitism and grafting.

**Fig. 2.**
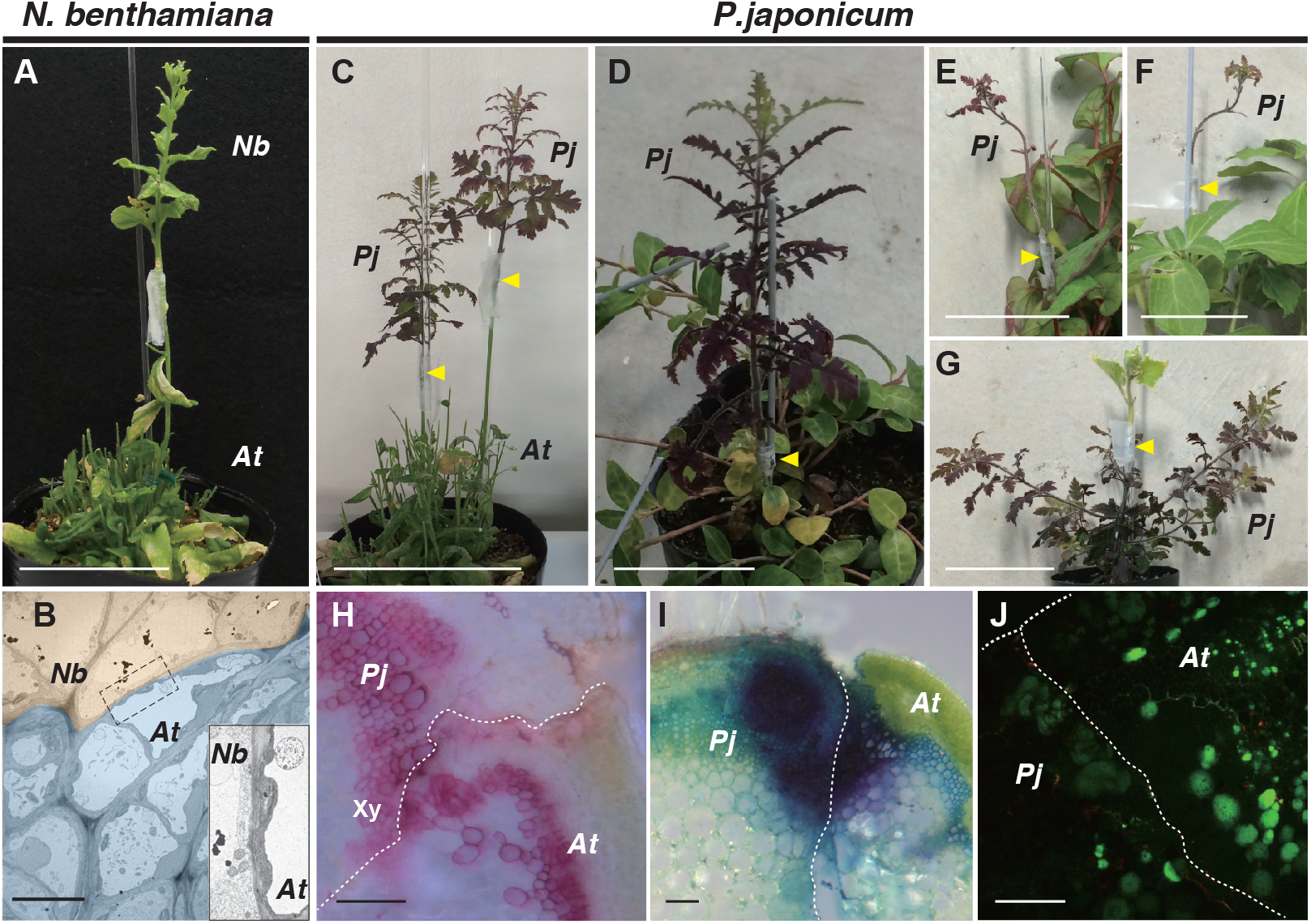
Heterospecific grafting of *N. benthamiana* and *P. japonicum*. (*A*) Grafting of *N. benthamiana* onto *Arabidopsis* at 28 days after grafting (DAG). (*B*) Transmission electron micrograph of the graft junction between *N. benthamiana* (pink) and *Arabidopsis* (blue). Inset represents a magnified image of the area of the dashed rectangle. (*C*–*G*) Grafting of *P. japonicum* with diverse plant species; grafts of *P. japonicum* as the scion onto stems of *Arabidopsis* at 28 DAG (*C*), *Vinca major* at 52 DAG (*D*), *Houttuynia cordata* at 30 DAG, and (*E*) *Pachysandra terminalis* at 30 DAG, and (*F*) graft of *Cucumis sativus* as the scion onto *P. japonicum* stock at 30 DAG (*G*). Arrowheads indicate grafting points. (*H*) Cross-section of the graft junction of *P. japonicum*/*Arabidopsis*. Phloroglucinol staining showing xylem formation in the graft region (Xy). (*I*) Cross-section of the graft junction of *P. japonicum*/*Arabidopsis* stained with toluidine blue applied to the *Arabidopsis* stock at 14 DAG. Detection of dye transport from *Arabidopsis* to *P. japonicum* demonstrated establishment of apoplastic transport. (*J*) Symplasmic transport establishment was confirmed using carboxyfluorescein (CF). CF was applied in the diacetate form to the leaves of the *Arabidopsis* stock and a cross-section of the graft junction of *P. japonicum*/*Arabidopsis* was observed. The CF fluorescence was detected in *P. japonicum* tissues. *Pj*, *P. japonicum*; *At*, *Arabidopsis.* Scale bars, 5 cm (*A*, *C*–*G*), 1 μm (*B*), and 100 μm (*H*–*J*).

**Table 1.**
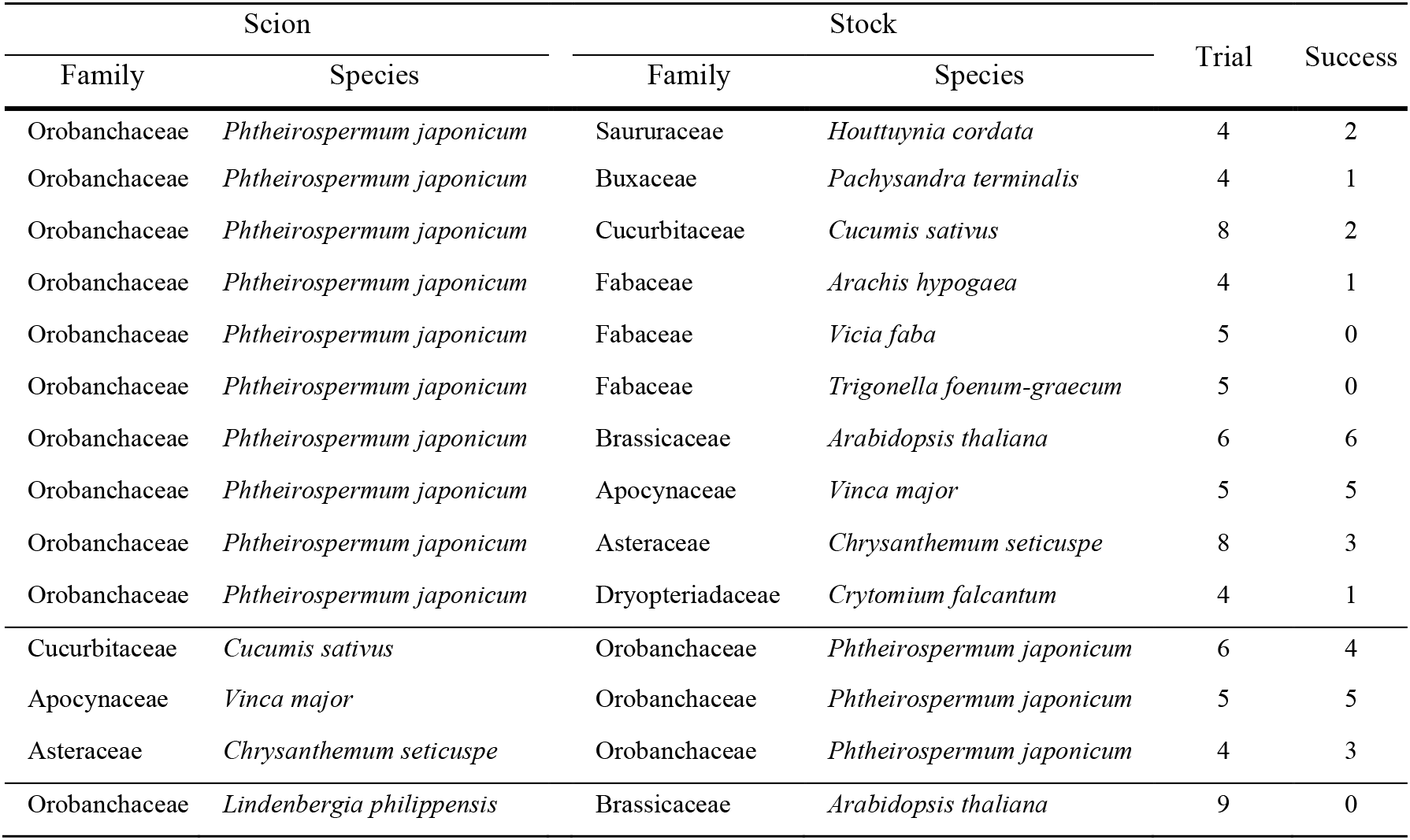
Grafting experiments performed with a root-parasitic plant, *Phtheirospermum japonicum*.

### Transcriptome analyses of parasitism and grafting

To investigate molecular events involved in cell–cell adhesion between *P. japonicum* and the host plant, we analyzed the transcriptome for *P. japonicum*–*Arabidopsis* parasitism and *P. japonicum*/*Arabidopsis* grafting. For this purpose, sequential samples of haustorial infection sites at the roots from 1 to 7 days after inoculation (DAI) and of grafted region on the stems from 1 to 14 days after grafting (DAG) were collected and subjected to RNA sequencing (RNA-Seq) analysis. Preliminary analysis of these data sets by principal component analysis (PCA) indicated that the transcriptomes of *P. japonicum*–*Arabidopsis* parasitism of the root and *P. japonicum*/*Arabidopsis* grafting onto the shoot differed substantially. The two transcriptomes showed a relatively similar distribution on PC1, but were widely separated on PC2 (Fig. 3*A*). Hierarchical clustering also indicated that gene expression patterns during parasitism and grafting were different over all (Fig. *3B*). Numerous genes were highly upregulated during parasitism or grafting, including some genes previously known to be associated with wound healing processes, such as auxin action, procambial activity, and vascular formation (Fig. 3 *B* and *C*) (18, 25, 26). However, the expression of many genes was distinct between parasitism and grafting. For example, *PIN1*, which encodes an auxin efflux transporter, *cyclin B1;2*, a cell-cycle regulator, *PLL1*, involved in maintenance of the procambium, *VND7*, a NAC domain transcriptional factor that induces transdifferentiation of various cells into protoxylem vessel elements, and *OPS*, a regulator of phloem differentiation, were all upregulated in a similar manner in both parasitism and grafting. In contrast, *IAA1*, which encodes an auxin-induced regulator, *ANAC071*, a transcriptional factor involved in tissue reunion after wounding, and *LBD16* and *LBD29*, LOB domain proteins involved in lateral root development, were upregulated only in grafting but not in parasitism. Conversely, *WOX4*, which encodes a WUSCHEL-related homeobox protein maintaining cambium activity, and *TMO6*, which encodes a Dof-type transcriptional factor for vascular development, were upregulated only in parasitism but not in grafting. The expression levels of *IAA18*, which encodes an auxin-induced regulator, *PLT3* and *PLT5*, which encode AP2-domain transcription factors involved in root stem cell identity, *CESA4*, which encodes cellulose synthase involved in secondary cell wall biosynthesis, and *ALF4*, involved in lateral roots initiation, showed little change or tended to decrease in both parasitism and grafting.

**Fig. 3.**
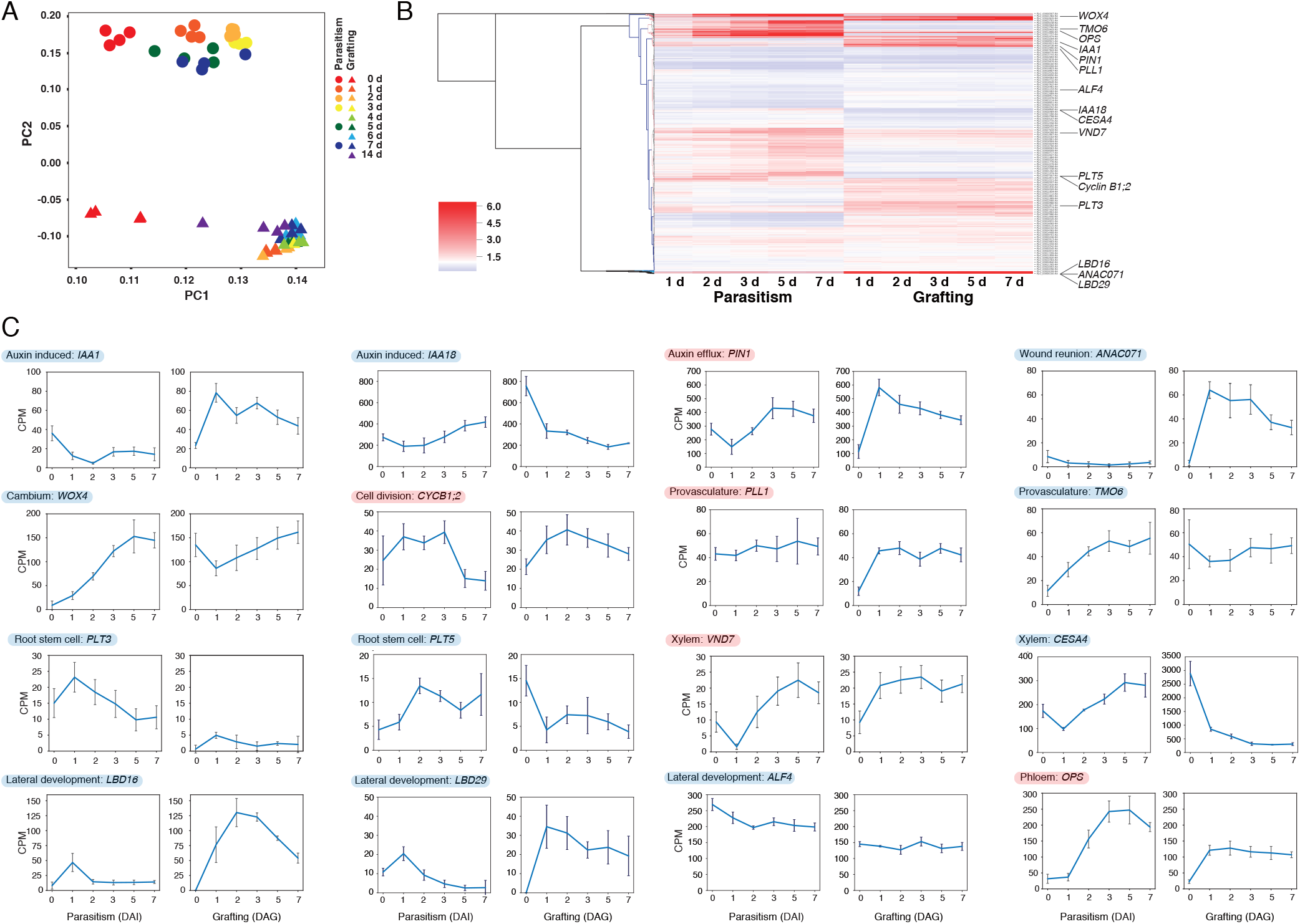
Transcriptomic analysis of parasitism and grafting between *P. japonicum* and *Arabidopsis*. (*A*) Transcriptomic analysis was performed using RNA samples of the *P. japonicum* infected site and the *P. japonicum* graft site. For parasitism, total RNA was extracted before and 1, 2, 3, 5, and 7 days after infection. For grafting, total RNA was extracted before and 1, 2, 3, 4, 5, 6, 7, and 14 days after grafting (with four biological replicates for each time point). Principal component analysis was performed from the obtained expression profile. Triangles and circles represent parasitism and grafting, respectively. PC, principal component. (*B*) Hierarchical clustering using Euclidean distance and Ward’s minimum variance method over ratio of RNA-seq data from five time points for *P. japonicum*–*Arabidopsis* parasitism and *P. japonicum*/*Arabidopsis* grafting against intact plants. Genes for which association with parasitism and grafting has been reported in previous studies are marked. Using the cDNA sequence of represents genes that behaved similarly in parasitism and grafting, and blue background indicates genes that behaved differently.

To identify common molecular events, we generated clusters for parasitism and grafting by multivariate analysis using self-organized map clustering, and compared clusters that included genes upregulated during tissue adhesion in parasitism and grafting (Fig. 4, *SI Appendix*, Fig. S1). During *P. japonicum* parasitization of the host root, tissue adhesion between *P. japonicum* and the host occurred around 1 to 2 DAI, and then a xylem bridge connecting a *P. japonicum* root vessel and a host vessel was formed at 3 DAI (Fig. 4*A*). In contrast, histological observation showed that tissue adhesion between the scion and stock during grafting occurred about 3 DAG (Fig. 4*C*). We focused three clusters with distinct expression patterns (Fig. 4*B*). The first one includes genes upregulated during the tissue adhesion stage starting about 1 DAI or 1 DAG (Node 09 for parasitism, Node 08 for grafting). The second one contains genes upregulated along the time (Node 05 for parasitism and Node 08 for grafting). The third one includes genes peaked around 2 DAI or 3 DAG (Node 08 for parasitism and Node 11 for grafting). Gene ontology (GO) enrichment analysis revealed many of the enriched GO terms were common to parasitism and grafting (*SI Appendix*, Datasets S3–S5). Especially in the focused three clusters, many of the enriched GO terms were associated with cell division, such as DNA replication, cytoplasm, and RNA metabolism, which reflected the occurrence of active cell proliferation in the haustorium and the graft interface. Importantly, these clusters also included enriched GO terms associated with the cell wall common to parasitism and grafting (Fig. 4*B*, *SI Appendix*, Datasets S3–S5). To identify genes specifically associated with tissue adhesion, we further compared these data with transcriptome data from grafting between *Nicotiana* and *Arabidopsis* (20) (Fig. 5*A*). We identified 9 genes commonly upregulated during tissue adhesion period in the three distinct experiments, including genes associated with cell division, such as cyclin D, and cell wall-related genes, such as glycosyl hydrolase (Fig. 5*A*). One of the identified glycosyl hydrolases belonging to the *Glycosyl hydrolase 9B* (*GH9B*) family, which includes genes encoding cellulases in plants (27, 28). Interestingly, among the *GH9B* family, a member of *GH9B3* clade was recently shown to be crucial for cell–cell adhesion in *Nicotiana* interfamily grafting (20).

**Fig. 4.**
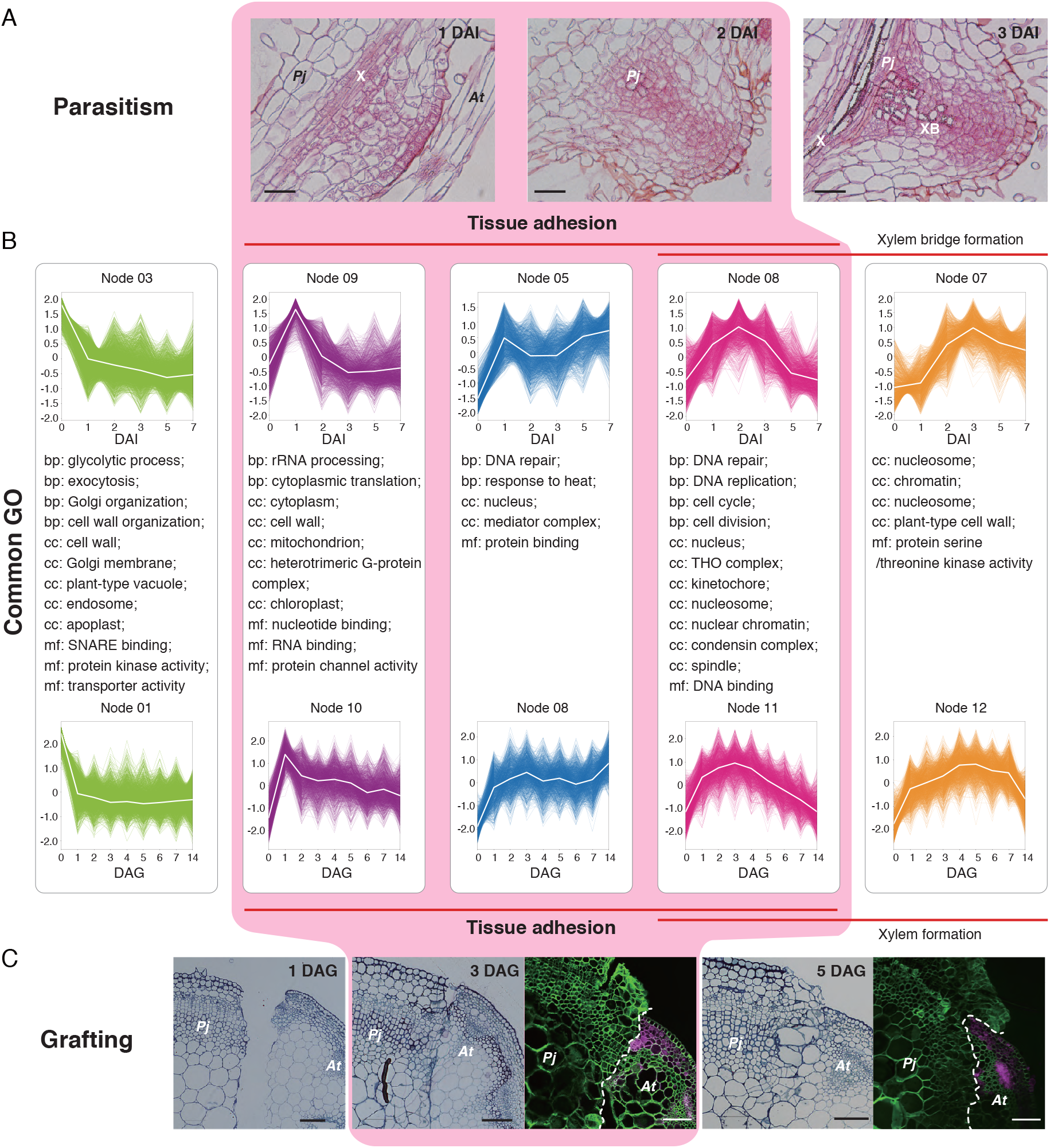
Comparison of self-organizing map (SOM) clusters associated with the tissue adhesion stage during parasitism and grafting. (*A*) Tissue sections of the parasitized site are shown. X represents the xylem of *P. japonicum* and XB represents the xylem bridge in the haustorium. (*B*) SOM clusters with similar patterns in parasitism (top) and grafting (bottom) are shown. Enriched gene ontology (GO) terms common to parasitism and grafting are listed for the clusters. bp, cc, and mf represent the GO subcategories biological process, cellular components, and molecular function, respectively. GO terms common to three or more pairs of cluster were excluded. For mf and bp, only GO terms with *P* < 0.01 were targeted. (*C*) Tissue sections of the graft junction are shown. Fluorescence images of the graft junction are also shown where *P. japonicum* was grafted onto *Arabidopsis* harboring *RPS5a::LTI6b-tdTomato*. Green indicates the cell wall, magenta indicates tdTomato fluorescence. Scale bars, 50 μm (*A*), and 100 μm (*C*).

**Fig. 5.**
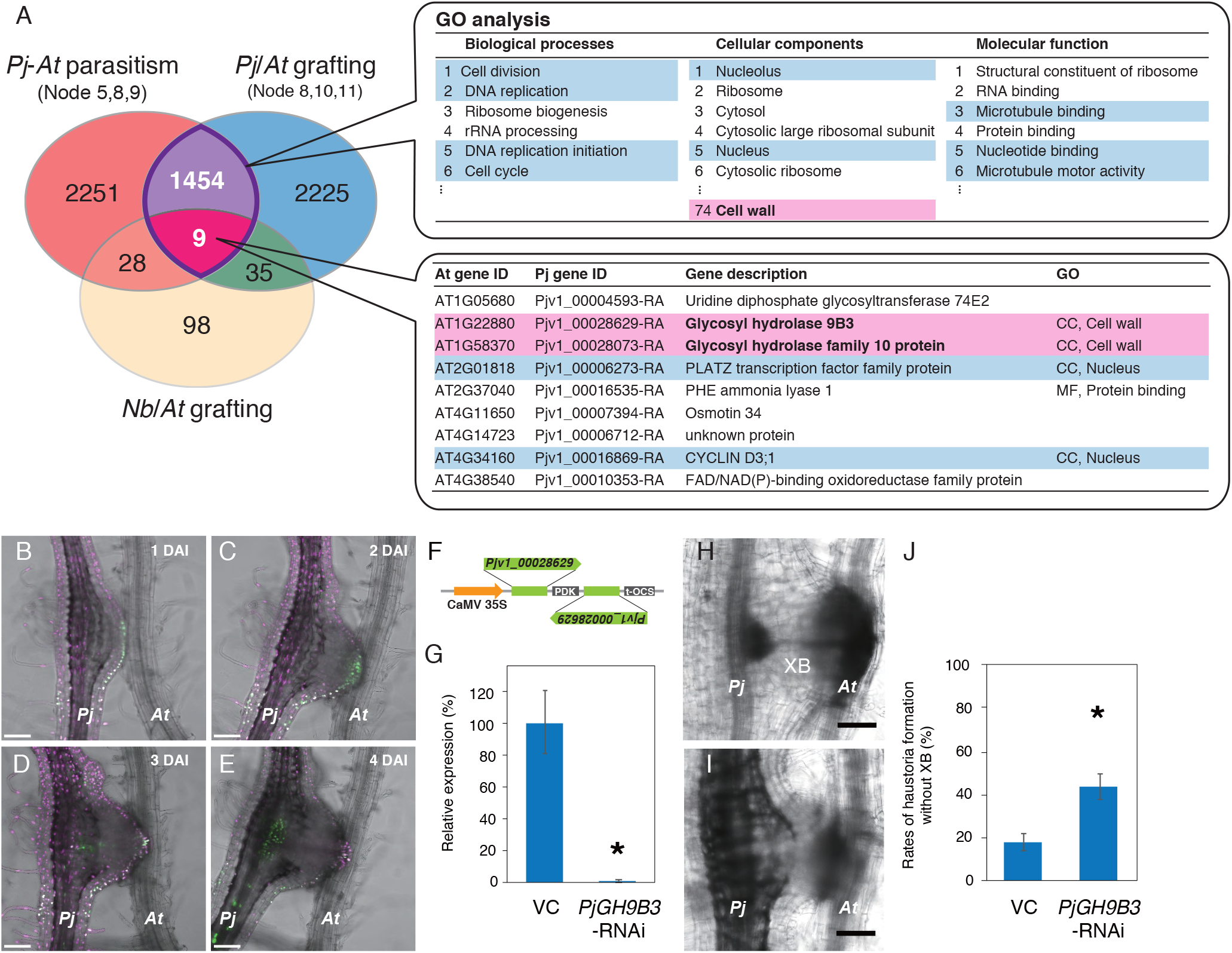
Involvement of *Glycosyl hydrolase 9B3* in establishment of parasitism of *P. japonicum*. (*A*) Extraction of factors important for tissue adhesion in parasitism and grafting. Nodes 5, 8, and 9 were merged from the self-organizing map (SOM) cluster of the *P. japonicum* parasitism transcriptome, and nodes 8, 10, and 11 were merged from the SOM cluster of the *P. japonicum* grafting transcriptome. Venn diagrams of the three gene populations were plotted, together with 170 *Arabidopsis* genes annotated by 189 *N. benthamiana* genes for which expression was elevated in the early stage of interfamily grafting of *N. benthamiana* and *Arabidopsis* (20). From the results of gene ontology (GO) analysis of the 1463 genes detected for both *P. japonicum* parasitism and grafting, a portion of the GO terms categorized as ‘biological processes (BP)’, ‘cellular components (CC)’ and ‘molecular functions (MF)’ are shown. Nine genes common to the three gene datasets are listed. GO terms and genes potentially associated with ‘cell division’ and ‘cell wall’ are marked in red and blue, respectively. (*B*–*E*) Expression patterns of *PjGH9B3* promoter::*GFP* 1, 2, 3, and 4 days after infection (DAI). (*F*) RNAi targeting sequence for *PjGH9B3* and the construct. (*G*) Relative expression levels of *PjGH9B3* at *PjUBC2* was used as a reference gene. Asterisk indicates statistical significance (Welch’s *t*-test, *P* < 0.05). VC, vector control. (*H, I*) Representative images of the haustoria that did (*H*) and did not (*I*) form a xylem bridge (XB) at 4 DAI. (*J*) Ratio of the haustoria that did not form a XB (mean ± SE of four replicates, *n* = 4–26). VC, vector control. *Pj*, *P. japonicum*; *At*, *Arabidopsis*; XB, xylem bridge. Scale bars, 100 μm.

### *PjGH9B*3 is essential for *P. japonicum* parasitism

As cell walls locating at the interface between *P. japonicum* and *Arabidopsis* were partially digested (Fig. 1 *E* and *F*), we decided to analyze function of the *GH9B3* in parasitism. We reconstructed a phylogenetic tree for the *GH9B* gene family for *Arabidopsis*, *P. japonicum*, and *S. hermonthica*, as well as *L. philippensis*, a non-parasitic Orobanchaceae species (29) (*SI Appendix*, Fig. S2*A*). In the phylogenetic tree, five and four genes from *P. japonicum* and *S. hermonthica* are found in the *GH9B3* clade, respectively, while only two and one *GH9B3* genes are present in *Arabidopsis* and *L. philippensis*, respectively. The *LiPhGnB1_1726*, a member of the *L. philippensis GH9B3* clade was upregulated at 1 DAG, but such upregulation was observed in both compatible and incompatible graft combinations using other plant species; soybean (*Glycine max*), morning glory (*Ipomoea nil*) and *Arabidopsis* (20). However, expression did not continue to increase subsequently, as this is often the case for incompatible graft combinations (20) (*SI Appendix*, Fig. S2*C*). By contrast, *Pjv1_00028629-RA*, the most similar *P. japonicum* homolog of *NbGH9B3* gene that was associated with grafting, was upregulated at 1 DAG and gradually increased until 7 DAG in grafting, as well as 7 DAI in parasitism (*SI Appendix*, Fig. S2*C*). Similarly, in *S. hermonthica*, the corresponding homolog *Sh14Contig_25152* was upregulated at 1 DAI with a peak at 3 DAI (17) (*SI Appendix*, Fig. S2*C*). These data suggest that upregulation of *GH9B3* homologs in parasitic plants at the site of infection and interfamily grafting is conserved in parasitic Orobanchaceae plants.

*Pjv1_00028629-RA* is upregulated at the early phase of infection and is the most closely related to homologs from *Nicotiana* and *Arabidopsis GH9B3* genes. Therefore, we investigated the function of *Pjv1_00028629-RA*, called *PjGH9B3*. First, temporal and spacial expression patterns of *PjGH9B3* were examined using a promoter-GFP construct. *PjGH9B3* promoter activity was detected at the cell periphery of the haustorium attaching to the host at 1 to 2 DAI (Fig. 5 *B* *and C*), while the signal was later shifted to the inside of haustorium at 3 to 4 DAI (Fig. 5 *D* and *E*). These expression patterns indicated that *PjGH9B3* functions at the interface opposing the host root tissue at an early adhesion stage and xylem formation at the later stage. *PjGH9B3* knockdown experiments by RNA interference (RNAi) system revealed that significantly fewer successful xylem connections with host, compared to the control (Fig. 5 *F–J*). Reduction of *PjGH9B3* expression was confirmed in hairy roots using quantitative reverse-transcription PCR (RT-qPCR) (Fig. 5*G*). These data indicate that *PjGH9B3* positively regulates infection processes in *P. japonicum*.

## Discussion

Parasitic invasion by haustorium is established through the disruption of host cell wall barriers. On the other hand, similar modification of the cell wall is also observed in the graft, which is an artificial tissue connection. In general, grafting is successful between closely related plants, such as members of the same species, genus, and family, but not between genetically distant plants, such as members of different families, with a few exceptions, such as *Nicotiana*, which we previously reported (10, 11, 20, 30, 31). We speculated that there might be a commonality between the capability of these parasitic plants and the capability to realize heterografting. In fact, interfamily grafting of parasitic plants resulted in successful with multiple combinations (Fig. 2 *C*–*G*, Table 1). We grafted *Phtheirospermum japonicum* as a scion mainly because cell proliferation at the graft surface on scion side is generally more active. In the case using Fabaceae as a graft partner, one species succeeded to graft but two species did not, indicating that appearance of graft incompatibility is independent to phylogenetic relation, rather more likely depends on each plant character or each combination. Graft capability of *P*. *japonicum* was also observed when *P*. *japonicum* was grafted as a stock (Fig. 2 *C*–*G*, Table 1). Furthermore, *L. philippensis*, a non-parasitic Orobanchaceae species, failed to establish interfamily grafting. These facts imply that parasitic plants may have acquired a mechanism to reconstruct the cell walls of plant tissues different from themselves in the evolution of parasitism.

Plant cells form primary and secondary cell walls. In general, primary cell walls are synthesized in cells of growing tissues and are composed predominantly of cellulose, pectic polysaccharides, and hemicelluloses such as xyloglucans. Secondary cell walls are formed in mature cells and are composed of cellulose, xylans, and lignin (27). Parasitic plants express cell wall modifying enzymes at the contact site and loosen cell walls at the host– parasite interface to invade the host tissues. *P. japonicum* expresses β-1,4-glucanases of the GH9B family, which show secretory signal peptides at the periphery of the haustorium at 1– 2 DAI (Fig. 5). The GH9B gene family is highly conserved in plants (28). Previous studies of microbial GH9 proteins have shown that these enzymes generally cleave the β-1,4-linkages of the glycosyl backbones of amorphous cellulose (32). *CELLULASE 3* (*CEL3*) and *CEL5* are GH9B3 clade homologs in *Arabidopsis* that show cellulase activities and are considered to be involved in cell loosening and expansion in lateral roots and detachment of root cap cells (33–35). We recently showed that GH9B3 clade genes play roles in cell–cell adhesion in graft junctions of *Nicotiana*, and *Arabidopsis* (20). Thus, parasitic plants activate conserved cell wall degrading enzymes that target cellulose, the predominant cell wall polysaccharide of various cell types in plants. This phenomenon may partially account for the compatibility in tissue adhesion of parasitic plants with diverse host plant species.

Gene duplication is involved in the evolution of parasitism. A previous study showed that duplication events occurred in more than half of the parasitic genes of Orobanchaceae species, which comprise a large number of genes annotated with GO terms associated with cell wall modifying enzymes and peptidase activity (12, 18). We identified five genes from *P. japonicum* that are classified in the *GH9B3* clade. The *GH9B8* clade also includes four genes from *P. japonicum* and one gene shows slight upregulation during parasitism. In contrast, no notable expansion was observed for other genes of the GH9 family. In addition, *GH9B3* of nonparasitic *L. philippensis* comprised only one gene, whereas the parasitic *S. hermonthica* showed duplication of *GH9B3* genes similar to that of *P. japonicum*. Gene duplication observed in the KAI2 strigolactone receptor family in *S. hermonthica* is also considered to be a driver for successful parasitism (4). The number of *GH9B3* clade genes in *P. japonicum* is not notably higher compared with KAI2 duplication in *S. hermonthica*. On the other hand, gene duplication in the Orobanchaceae predominantly occurred before its divergence from nonparasitic plants and diversification in the regulation of the duplicated genes may also have contributed to the evolution of parasitism (12). Although the amino acid sequences were highly conserved, we assumed that gene expression patterns can be varied by increase gene copy. Although all homologous *GH9B3* genes in the Orobanchaceae parasitic plants we examined encode functional proteins, their expression patterns were temporally varied (Fig. 5 *B*–*E*, *SI Appendix*, Fig. 2 *C* and *D*). Taken together, we assume that duplication of *GH9B3* and the resulting diversification in spatiotemporal expression were both important factors in the evolution of parasitism.

We narrowed the genes associated with tissue adhesion among distant species (Figs. 4 and 5) and identified a role in parasitism for one of candidate genes using *P. japonicum* system (Fig. 5). Additional important components are likely to facilitate cell wall digestion and accomplish tissue adhesion. The dataset accumulated in the present study provides a foundation to identify such components involved in parasitic infection and/or grafting. *P. japonicum* is a useful model system because the seeds can germinate in the absence of a host plant, obtain transformants from hairy roots, parasitize the host, and can be grafted to the host plant species (15, 21, 22, 36) (Figs. 1 and 2). Improved knowledge of the mechanisms responsible for parasitism could be applicable to suppress yield losses in crop cultivation caused by parasitic plants. The present study may provide an option, which might be especially effective against hemiparasites, to decrease parasitization by parasitic plants after germination by inhibiting the activities of secreted endo-β-1,4-glucanases. Several mono- and polysaccharides are reported to be inhibitors of cellulase (37). Hence, a knowledge-based defense approach would further enhance crop security.

## Materials and Methods

### Plant materials

For the grafting experiments, *Phtheirospermum japonicum* and *Arabidopsis thaliana* ecotype Columbia (Col-0) seeds were directly surface-sown on soil. Plants were grown at 22°C under continuous light. Infection assay was performed as described previously (22). Briefly, *Arabidopsis* seeds were surface-sterilized with 70% ethanol for 10 min, washed three times with sterile water, incubated at 4°C for 2 days, and sown on half-strength Murashige and Skoog (1/2 MS) medium (0.8% agar, 1% sucrose, pH 5.8). Seedlings were grown vertically at 22°C under long-day (LD) conditions (16 h light/8 h dark). *P. japonicum* seeds were surface-sterilized with 10% (v/v) commercial bleach solution (Kao, Tokyo, Japan) for 5 min, rinsed three times with sterilized water, and sown on 1/2 MS agar medium (0.8% agar, 1% sucrose). The agar plates were incubated at 4°C in the dark overnight, then incubated at 25°C under LD. Plants for the infection assay and the transformation experiment were cultured vertically and horizontally, respectively.

### Grafting

For grafting of *P. japonicum*/*Arabidopsis*, 1- to 2-month-old *P. japonicum* and 1-month-old *Arabidopsis* plants were used. Wedge grafting was performed on the epicotyls, stems, petioles, or peduncles. For stock preparation, stems (or other organs) were cut with a 2–3 cm slit on the top. For scion preparation, the stem (around 7~10 cm from the tip of the sttem) was cut and trimmed into a V-shape. The scion was inserted into the slit of the stock and wrapped with parafilm to maintain close contact. A plastic bar was set along the stock and the scion to support the scion and the graft. The entire scion was covered with a plastic bag, which had been sprayed inside with water beforehand. Grafted plants were grown for 7 days in an incubator at 27°C under continuous light (~30 μmol m^−2^ s^−1^). After this period, the plastic bags were partly opened by cutting the bags and making holes for acclimation. The next day, the plastic bags were removed and the grafted plants were transferred to a plant growth room at 22°C under continuous light conditions (~80 μmol m^−2^ s^−1^). All other plant materials for stem grafting used in this study are listed in Table 1.

### Microscopy

All the chemicals for staining were purchased from Sigma-Aldrich Co., Tokyo, Japan unless otherwise stated. Preparation of resin sections and observation by a transmission electron microscope were performed as previously described (20). To capture images of infection tissues or hand-cut grafted regions, a stereomicroscope (SZ61, Olympus, Tokyo, Japan) equipped with an on-axis zoom microscope (Axio Zoom.V16, Zeiss, Göttingen, Germany). To observe xylem tissues, phloroglucinol staining (Wiesner reaction) was performed by applying 18 μL of 1% phloroglucinol in 70% ethanol followed by the addition of 100 μL of 5 N hydrogen chloride to the section samples. To determine apoplastic transport, the stems of *Arabidopsis* stocks were cut and the cut edge was soaked in 0.5% toluidine blue solution for 4 h to overnight. To determine symplasmic transport, cut leaves of the *Arabidopsis* host or stock were treated with 0.01% 5(6)-carboxyfluorescein (CF) diacetate. The CF fluorescence images were acquired by a fluorescence imaging stereomicroscope or observing emissions in the 490–530 nm range with excitation at 488 nm with a confocal laser scanning microscopy (LSM780, Zeiss). To examine graft junctions and visualize tissue adhesion, grafting was performed between a *P. japonicum* scion and a transgenic *Arabidopsis* stock, *RPS5A*::*LTI6b*-*tdTomato*, that ubiquitously expresses a plasma membrane-localized fluorescent protein (38). Hand-cut cross-sections of the grafted stem regions were stained with 0.001% calcofluor white, which stains cellulose in plant cell walls. The fluorescence of tdTomato or calcofluor white was detected using a confocal laser scanning microscope (LSM710, Zeiss). A 555 nm laser and collecting emission spectrum of 560–600 nm were used for tdTomato and a 405 nm excitation laser and collecting emission spectrum of 420– 475 nm were used for calcofluor white. The GFP or mCherry fluorescence images were acquired as described previously (21).

### Transcriptome analysis

RNA sequencing was conducted for sequential samples of haustorial infection sites in the roots and of grafted region on the stems as described previously (20, 23, 41). For parasitism and grafting samples between *P. japonicum* and *Arabidopsis*, the sequence reads were mapped on the genome assembly using TopHat version 2.1.0 with default parameters (http://ccb.jhu.edu/software/tophat/). Uniquely mapped reads are counted using HTSeq (https://htseq.readthedocs.io/en/master/), and high homology genes that show more than 10 total reads of *P. japonicum* or *Arabidopsis* samples mapped on *A. thaliana* or *P. japonicum* genome, respectively, were removed. These mapped reads were normalized using the Bioconductor package EdgeR ver. 3.4.2 (https://bioconductor.org/packages/release/bioc/html/edgeR.html) with the trimmed mean of M-values method where reads were filtered like that there was at least one count per million (CPM) in at least half of the samples. For the sample of grafting between *L. philippensis, Nicotiana* and *Arabidopsis*, the sequence reads were mapped on the genome assembly using HISAT2 version 2.1.0 (http://daehwankimlab.github.io/hisat2/). Gene expression levels, expressed as fragments per kilobase of transcript per million fragments mapped (FPKM), were estimated using Cufflinks version 2.1.1 (http://cole-trapnell-lab.github.io/cufflinks/). The reference sequence used for mapping and the annotation file used were as follows: *P. japonicum* PjScaffold_ver1; *N. benthamiana* draft genome sequence v1.0.1, https://btiscience.org/our-research/research-facilities/research-resources/nicotiana-benthamiana/; *A. thaliana* TAIR10 genome release, https://www.arabidopsis.org; and *L. philippensis* LiPhGnB1, http://ppgp.huck.psu.edu. Gene ontology enrichment analysis was performed with DAVID (https://david.ncifcrf.gov) using *Arabidopsis* gene IDs. Transcriptome data of parasitism and grafting were used for principle component analysis (PCA) to compare the differences between samples. The Python modules including numpy, pandas, matplotlib, seaborn and Scikit-learn were used for PCA and hierarchical cluster classification. For SOM clustering, the parasitism was classified into nine clusters and the grafting was classified into 12 clusters. After clusters with similar patterns were paired, in three clusters expression increased when cell adhesion occurred in both the parasitism and the grafting, and the clusters located before and after those were presented. The RNA-Seq data are available from the DNA Data Bank of Japan (DDBJ; http://www.ddbj.nig.ac.jp/).

### Plasmid construction

We used Golden Gate modular cloning to construct a vector to examine the expression pattern of *PjGH9B3* during infection (40). The *PjGH9B3* promoter region (1899 bp upstream of the ATG start codon) was cloned into the vector pAGM1311 as two fragments and then combined into the vector pICH41295 to generate the promoter module. This module was assembled into the vector pICH47751 containing 3xVenus-NLS and AtHSP18.2 terminator (21), then subsequently further assembled into pAGM1311 with *pPjACT::3xmCherry-NLS* (21). For RNAi experiments, we used the pHG8-YFP vector (41). Target sequences were PCR-amplified from *P. japonicum* genomic DNA and cloned into the pENTR vector (Thermo Fisher Scientific, Waltham, MA, USA), then transferred into the pHG8-YFP vector by the Gateway reaction using LR Clonase II Plus enzyme (Thermo Fisher Scientific). All primers used in this paper are listed in Dataset S10.

### *Phtheirospermum japonicum* transformation

*P. japonicum* transformation was performed as previously described (36). Six-day-old *P. japonicum* seedlings were sonicated and then vacuumed for 10 s and 5 min, respectively, in an aqueous suspension of *Agrobacterium rhizogenes* strain AR1193. After incubation in freshly prepared Gamborg’s B5 medium (0.8% agar, 1% sucrose, 450 μM acetosyringone) at 22°C in the dark for 2 days, seedlings were incubated in Gamborg’s B5 medium (0.8% agar, 1% sucrose, 300 μg/ml cefotaxime) at 25°C under LD.

### Parasitization assay and RT-qPCR

Ten-day-old *P. japonicum* seedlings were transferred from 1/2 MS medium agar plates to 0.7% agar plates and incubated at 25°C under LD for 2 days. Seven-day-old *Arabidopsis* seedlings were placed next to *P. japonicum* seedlings so that *P. japonicum* root tips could physically contact the *Arabidopsis* roots. For RNAi experiments, fresh elongated hairy roots were transferred to 0.7% agar plates and incubated at 25°C under LD for 2 days. YFP- or mCherry-positive hairy roots were placed between thin agar layers (0.7%) and glass slides in glass-bottom dishes (Iwaki, Haibara, Japan), and incubated at 25°C under LD overnight. Seven-day-old *Arabidopsis* seedlings were placed next to *P. japonicum* seedlings underneath agar layers so that *P. japonicum* root tips could physically contact the *Arabidopsis* roots. Xylem bridge formation and the expression pattern of *PjGH9B3* were examined using a confocal microscope (Leica SP5). After phenotyping xylem bridge formation, haustorial tissues were dissected and snap frozen in liquid nitrogen. Total RNA was extracted using the RNeasy Plant Mini kit (Qiagen, Hilden, Germany) and cDNA was synthesized using ReverTraAce qPCR RT Master Mix (TOYOBO, Osaka, Japan). The RT-qPCR analyses were performed using a Stratagene mx3000p quantitative PCR system (with 50 cycles of 95°C for 30 s, 55°C for 60 s, and 72°C for 60 s) with the Thunderbird SYBR qPCR Mix (TOYOBO). *PjUBC2* was used as an internal control as previously described (22). The primer sequences used are shown in Dataset 10. All experiments were performed with three independent biological replicates and three technical replicates.

## Acknowledgements

We thank S. Yoshida (Nara Institute of Science and Technology, Japan) for providing information relevant to the *P. japonicum* genome. We thank T. Shinagawa, H. Fukada, A. Ishiwata, and A. Shibata for technical assistance. This work was supported by grants from the Japan Society for the Promotion of Science Grants-in-Aid for Scientific Research (18KT0040, 18H03950, 19H05361 to M.N. and 15H05959, 17H06172 to K.S.), the Cannon Foundation (R17-0070), and the Japan Science and Technology Agency (START15657559 and PRESTO15665754) to M.N.

