## Supplemental file for "Host-parasite tissue adhesion by a secreted type of β-1,4-glucanase in the parasitic plant *Phtheirospermum japonicum*"

**Dataset S1. Expression data for parasitism between *P. japonicum* and *Arabidopsis*.**
**Dataset S2. Expression data for grafting between *P. japonicum* and *Arabidopsis*.**
**Dataset S3. List of GO terms in six clusters shown in Fig. 4*B* (BP category, *P* < 0.01).**
**Dataset S4. List of GO terms in six clusters shown in Fig. 4*B* (CC category).**
**Dataset S5. List of GO terms in six clusters shown in Fig. 4*B* (MF category, *P* < 0.01)**
**Dataset S6. List of genes overlapping between parasitism and grafting of *P. japonicum* shown in Fig. 5*A*.**
**Dataset S7. List of GO terms in genes overlapping between parasitism and grafting of *P. japonicum* shown in Fig. 5*A* (BP category).**
**Dataset S8. List of GO terms in genes overlapping between parasitism and grafting of *P. japonicum* shown in Fig. 5*A* (CC category).**
**Dataset S9. List of GO terms in genes overlapping between parasitism and grafting of *P. japonicum* shown in Fig. 5*A* (MF category).**
**Dataset S10. Primers used in this study.**

**Fig. S1. Whole self-organizing map (SOM) clusters and gene ontology (GO) terms for parasitism and grafting between *Phtheirospermum japonicum* and *Striga hermonthica.***
**Fig. S2. *Glycosyl hydrolase 9B* gene family in Orobanchaceae.**
**Fig. S3. Model of parasitism by *Phtheirospermum japonicum*.**

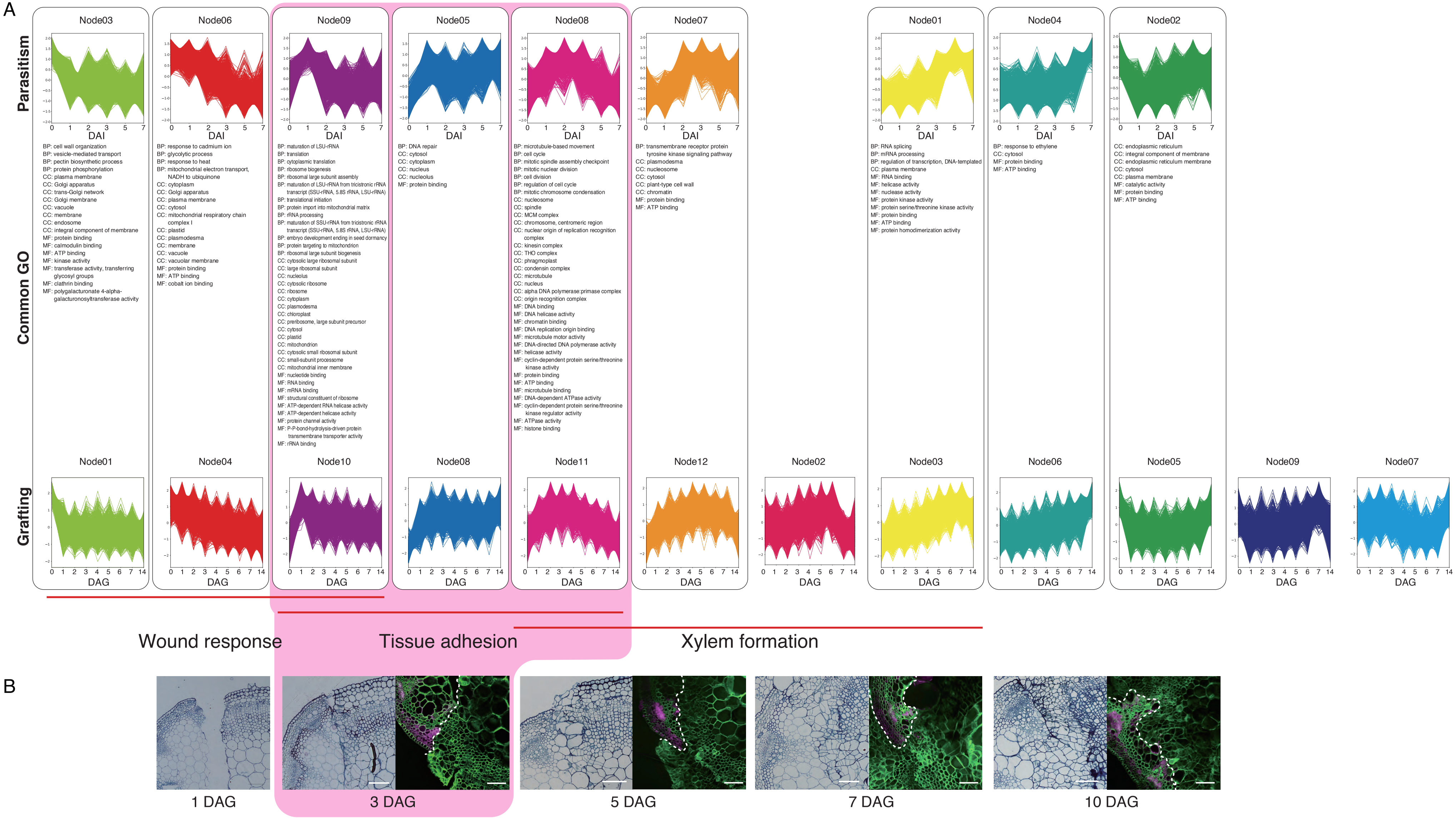


**Fig. S1.** Whole self-organizing map (SOM) clusters and gene ontology (GO) terms for parasitism and grafting between *Phtheirospermum japonicum* and *Arabidopsis.* (*A*) SOM clusters with similar patterns in parasitism (top) and grafting (bottom) are shown. GO terms commonly found in the parasitism and grafting were listed among clusters. BP, CC and MF stand for GO-subcategory of biological process, cellular components and molecular function, respectively. 20 GOs with a *P* value of 0.05 or less were selected in each category. (*B*) Tissue sections of the graft junction are shown. Fluorescence images of the graft junction are also shown where *P. japonicum* was grafted to *Arabidopsis RPS5a::LTI6b-tdTomato*. Green indicates the cell wall, magenta indicates tdTomato fluorescence. Scale bars, 100 µm.

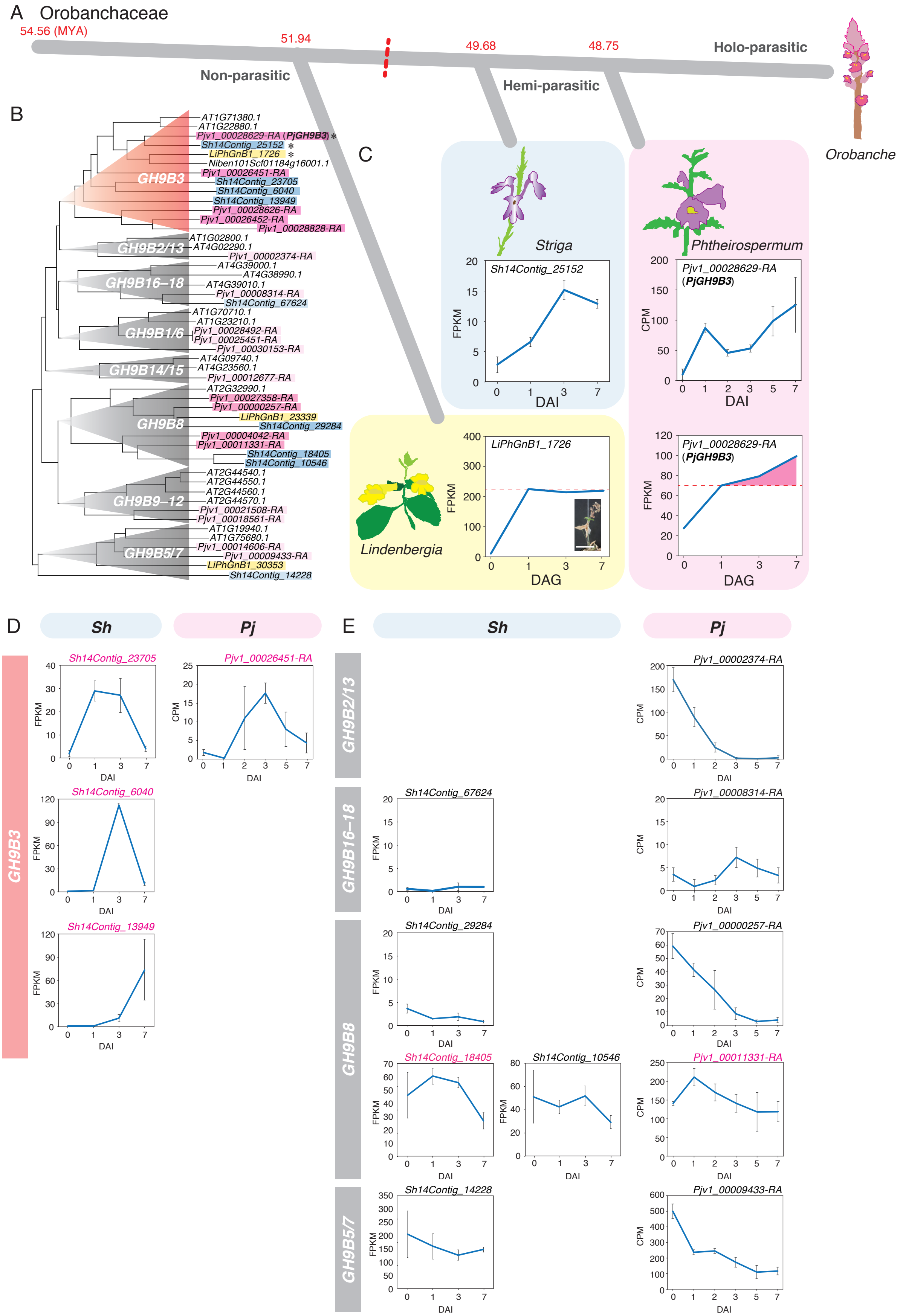


**Fig. S2.** *Glycosyl hydrolase 9B* gene family in Orobanchaceae. (*A*) Phylogeny of Orobanchaceae including non-, hemi-, a nd holoparasitic species. Estimated branching time points (million years ago) are indicated (29). (*B*) Phylogeny of *Glyco syl hydrolase 9B3* genes of *Lindenbergia philippensis*, *Phtheirospermum japonicum*, *Striga hermonthica*, *Arabidopsis*, and *Nicotiana benthamiana* reconstructed using deduced amino acid sequences. (*C*) Expression patterns of genes encoding the amino acid sequence closest to *NbGH9B3* and *AtGH9B3* in grafting for *L. philippensis* and *P. japonicum* or parasitism for *S. hermonthica* and *P. japonicum*. (*D*) Expression patterns of other genes of *S. hermonthica* belonging to the GH9B3 clade. (*E*) Expression patterns of other GH9B genes belonging to clades other than GH9B3 in *S. hermonthica* and *P. japonicum*. *Sh*, *S. hermonthica*; *Pj*, *P. japonicum*.

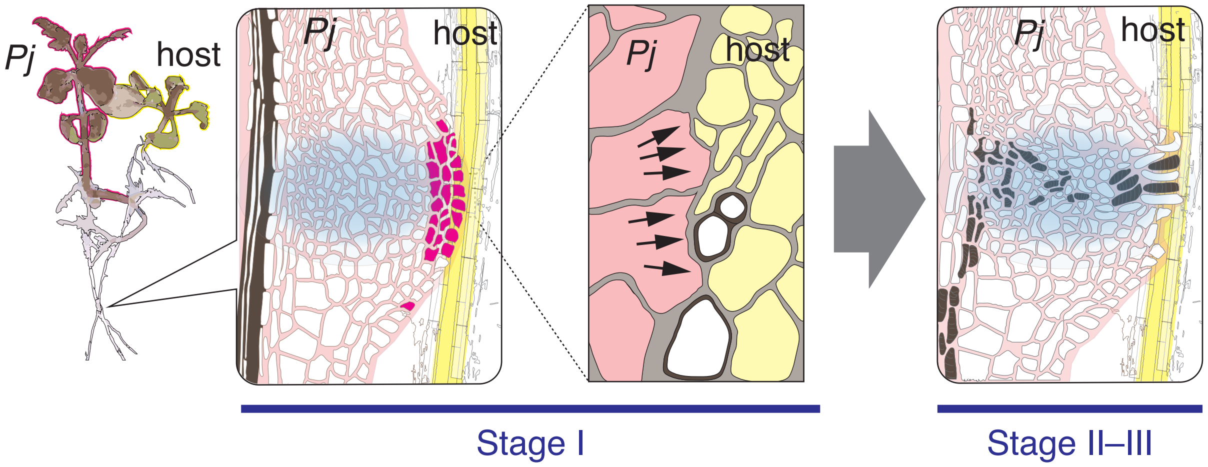


**Fig. S3.** Model of parasitism by *Phtheirospermum japonicum*. Upon detection of host plant roots, *P. japonicum* generates haustoria where cells are proliferated through cell division (colored in blue). *PjGH9B3* is induced at the periphery of the haustorium (colored in magenta) to facilitate tissue adhesion when in contact with the host plant root (stage I). Subsequently, a xylem bridge (colored in brown) is formed between *P. japonicum* and the host plant root (stages II–III). Arrows indicate decreased cell wall thickness at the interface between parasitic and host tissues. *Pj*, *P. japonicum.*
